# CEBPβ regulates myoblast proliferation and myogenic differentiation during human myogenesis and rescues defective differentiation in FSHD

**DOI:** 10.1101/2025.08.07.669099

**Authors:** Elise N. Engquist, Isabella Hofer, Johanna Pruller, Maryna Panamarova, Christopher R.S. Banerji, Peter S. Zammit

**Author notes:** Corresponding Author: Peter S. Zammit, King’s College London, Randall Centre for Cell and Molecular Biophysics, New Hunt’s House, Guy’s Campus, London SE1 1UL, UK.

## Abstract

CCAAT enhancer binding protein β (CEBPB) is a member of the bZIP transcription factor family and is expressed in many cell types. Whilst CEBPB has been shown to regulate the function of murine satellite cells, the resident skeletal muscle stem cell, however little is known of its role in human myogenesis and muscle disease. We report that CEBPB is expressed in proliferating human myoblasts but swiftly down-regulated upon myogenic differentiation. Interestingly, either over-expression or knock-down of CEBPB inhibit the proliferation rate of human myoblasts. Down-regulation of CEBPB upon myogenic differentiation is then necessary for subsequent myogenic differentiation and fusion into multinucleated myotubes. In the muscle disease context, CEBPB protein levels are increased in myoblasts derived from facioscapulohumeral muscular dystrophy (FSHD) patients. Knock-down of CEBPB in FSHD myoblasts improves myogenic differentiation. In summary, we show that CEBPB is a regulator of human myogenesis and altered CEBPB expression may contribute to defective myogenesis in FSHD.

## Introduction

CCAAT enhancer binding protein β (CEBPB) is a member of the bZIP transcription factor family. In humans, *CEBPB* is an intronless gene located on chromosome 20q13.13. *CEBPB* transcript encodes three protein isoforms, generated through translation initiation at different alternative start codons (1, 2). The canonical isoforms LAP* and LAP each contain a trans-activation domain, a regulatory domain, and a bZIP DNA-binding domain (1, 2). The truncated LIP isoform retains the DNA-binding and dimerization domains but lacks the trans-activation domain and part of the regulatory domain, and thus likely functions as a regulator of the transcriptional activity of LAP* and LAP (3).

CEBPB is expressed in a wide variety of cell types. CEBPB modulates differentiation and cytokine production in monocytes (4), inflammatory responses and energy metabolism in the liver, kidney, and vascular smooth muscle (5–8), and promotes adipocyte differentiation (9, 10). Circulating monocytes contain CEBPB protein (11), and *CEBPB* mRNA levels in blood are associated with muscle strength (12) as well as with response to exercise-induced muscle damage (13). CEBPB is also expressed by immune cells within skeletal muscle and is essential for the transition of macrophages from a pro-inflammatory role to one of supporting myogenesis during muscle regeneration (14, 15). Following muscle injury, macrophages with mutations in the murine *CEBPB* promoter still migrate to the damaged area and perform normal inflammatory responses but subsequently fail to activate anti-inflammatory programs, which ultimately contributes to severely impaired muscle regeneration (14, 15).

Post-natal skeletal myofibre growth, hypertrophy and maintenance, in addition to repair and regeneration, are performed by resident muscle stem cells termed satellite cells (16). Satellite cells are also recruited to repair muscle fibres in disease conditions, including in muscle wasting disorders such as muscular dystrophies. Importantly, *CEBPB* is expressed in quiescent murine satellite cells, with a role in maintaining the undifferentiated state (17, 18). *In vivo* models demonstrate that specific conditional deletion of *CEBPB* in satellite cells results in muscle fibre hypertrophy and improved muscle regeneration following acute injury (17).

However, satellite cells lacking *CEBPB* are gradually depleted over time, leading to impaired regeneration following multiple rounds of injury. This suggests that CEBPB is required for long-term maintenance and self-renewal of the murine myogenic stem cell pool (19). *CEBPB* is expressed both in primary murine myoblasts and C2C12 cells and is down-regulated upon myogenic differentiation (17, 20). Failure to down-regulate CEBPB blocks myogenic progression, while reduction of CEBPB improves myotube fusion and leads to aberrant differentiation in growth conditions (17, 20). PAX7 is a master regulator of post-natal satellite cell function, and CEBPB has been shown to bind directly to the *Pax7* promoter. This activates *PAX7* transcription as well as downstream quiescence-associated genes, and suppresses expression of myogenesis-associated genes such as *MyoD* and *Myogenin* (17, 18).

Our interest in CEBPB stems from our observation that *CEBPB* is mis-expressed in myogenic cells derived from patients with facioscapulohumeral muscular dystrophy (FSHD) (e.g. GSE102812). FSHD1 (OMIM 158900) and FSHD2 (OMIM 158901) are caused by genomic changes that result in epigenetic depression of D4Z4 units located on Chromosome 4, which ultimately leads to aberrant expression of the transcription factor *DUX4* on a permissive 4qA haplotype (21–23). However, the *DUX4* retrogene is restricted to old world primates (24), and so mouse models are limited.

While the role of CEBPB is well understood in mouse, little is known about CEBPB expression and function in human myogenesis, and there can be distinct differences between the role of regulators of myogenesis between mouse and human (e.g. (25)). Thus, we first needed to establish the role of CEBPB in human myoblasts. LAP*/LAP isoforms were present in proliferating human myoblasts, before being down-regulated during myogenic differentiation. Constitutive expression or knock-down demonstrated that CEBPB regulates human myoblast proliferation. Furthermore, CEBPB up-regulation largely prevents myogenic differentiation and fusion into multinucleated myotubes. Investigating FSHD, we found that *CEBPB* mRNA expression is altered in both patient biopsies and cell lines from multiple independent sources, and that FSHD myoblasts contain higher levels of CEBPB protein compared to controls. FSHD myoblasts often display a defect in myogenesis whereby they generate hypotrophic myotubes (26) but knocking down CEBPB in FSHD myoblasts improved their myogenic differentiation. Thus, mis-regulation of CEBPB may contribute to defective muscle regeneration in individuals affected by FSHD.

## Materials and Methods

### Human cell lines and culture reagents

Isogenic immortalized cell myoblast clones 54.12 (FSHD, 3 D4Z4 units) and 54.6 (control, 13 D4Z4 repeats) were obtained from Dr. Vincent Mouly (Institute of Myology, Paris, France) and were originally isolated from the biceps of a 54 year old mosaic male FSHD1 patient (27). 16A (FSHD) and 16U (sibling-matched control) were obtained from the UMMS Wellstone Centre for FSHD (Worcester, MA, USA), originally isolated from the biceps of a 56-year old female FSHD1 patient and her 60-year old unaffected sister (28).

Human myoblast lines were grown in a humidified incubator at 37 °C at 21% O_2_ and 5% CO_2_ in growth medium (GM) consisting of Skeletal Muscle Cell Growth Medium (Promocell, #C-23060) supplemented with 20% fetal bovine serum (FBS) (Gibco 10500064) and 50 μg/mL Gentamicin (Gibco, #157-50-060). To induce differentiation, myoblasts were grown to ∼90% confluency, washed twice in phosphate buffered saline (PBS), and placed in differentiation medium (DM) of DMEM Glutamax (Gibco, # 10569010) supplemented with 0.5% FBS, 10 μg/mL recombinant human insulin (Merck #I9278) and 50 μg/mL Gentamicin.

HEK293T cells used for lentiviral particle production were cultured in DMEM-GlutaMAX (ThermoFisher G31966-02), with 10% FBS and 1% penicillin-streptomycin (Sigma, #P0781).

### Protein Extraction and Western Blotting

Cells were harvested with Trypsin-EDTA (Sigma, #T4049) and incubated on ice for 30 min in RIPA buffer (ThermoFisher, #89900) containing Complete Protease Inhibitor Cocktail (Roche, #1187358000) and 50 mM sodium fluoride (Sigma, #67414), then centrifuged for 30 min at 4°C at 13000 rpm. Supernatant was collected and added 3:1 to a solution of Laemmli loading buffer (1610747, #Biorad) containing 10% β-mercaptoethanol. Protein samples were boiled for 5 min at 95°C and loaded in a 4-20% precast gel (BioRad, #4561094) alongside PageRuler Plus pre-stained ladder (ThermoFisher, #26620), separated by SDS-PAGE, and transferred to nitrocellulose membrane for Western Blot analysis.

Membranes were blocked in Tris-buffered saline with 0.1% Tween-20 detergent (TBS-T) + 5% milk for 1hr. Prior to primary antibody incubation, membranes were cut just below the 100 kDa line to separate vinculin (VCL) (117 kDa) from CEBPB protein isoforms (20-44 kDa). Respective membranes were incubated overnight at 4°C in either TBS-T + 5% milk + 1:10000 anti-vinculin primary antibody (Sigma, #V9131) or TBS-T + 5% BSA + 1:1000 anti-CEBPB primary antibody (Cell Signaling Technology, #90081), followed by secondary antibody incubation for 1 hr at room temperature (RT) in TBS-T + 5% milk + 1:5000 anti-rabbit-HRP (Sigma, #GENA934) or 1:5000 anti-mouse-HRP (Sigma, #NA931V) secondary antibodies. Membranes were developed in ECL substrate (Amersham, #RPN2232) and imaged on a ChemiDoc imaging system (BioRad, #17001401). Colorimetric images of ladders were collected immediately following chemiluminescent acquisition, and are shown alongside corresponding chemiluminescent images in figures. Quantification was performed in ImageJ, and relative density unit (RDU) for LAP*+LAP and LIP protein isoform expression was normalized to Vinculin (VCL).

### Generation of 54.6-iCEBPB and 16U-iCEBPB cell lines

A plasmid was designed in which the full-length CEBPB coding sequence is driven by the TRE promoter, and reverse tetracycline-controlled transactivator and mCherry are driven by ubiquitously active *SV40* and *CMV* promoters respectively and purchased from VectorBuilder (Supplementary Figure 1A). For lentiviral particle generation, the iCEBPB plasmid was incubated with pMDLg/pRRE, pRSV, PMD2.G plasmids (Plasmids #12251, #12253, #12259, Addgene) in OptiMEM (Gibco, #31985062) and transfected into HEK293T cells using Lipofectamine 3000 (Invitrogen, #L3000008). Medium was replaced after 24 hrs and viral supernatant was collected 48 hrs later, centrifuged at 1200 rpm for 3 min, and passed through a 0.45 μm filter. To generate 16U-iCEBPB and 54.6-iCEBPB cell lines, 16U/54.6 myoblasts were transduced with viral supernatant + 8 μg/mL Polybrene (Santa Cruz, #sc-134220) for 24 hrs, expanded for 3-5 passages, and FACS sorted for mCherry. CEBPB over-expression upon doxycycline induction was confirmed by immunofluorescence (Supplementary Figure 1B).

### siRNA knock-down of CEBPB

54.6 and 16U myoblast lines were transfected with 5 nM of either negative control siRNA (siNC) (Qiagen, #1027310; Negative Control siRNA) or anti-CEBPB siRNA (siCß) (Qiagen #1027416; FlexiTube GeneSolution GS1051, consisting of 4 siRNAs: SI02777292, SI03058062, SI03022341, SI0007364) in Lipofectamine RNAiMax (Invitrogen #13778075) per manufacturer’s instructions. For proliferating myoblasts, reverse transfection was performed at the time of cell seeding and medium was changed 24 hrs later. For knock-down during differentiation, siRNA was introduced with differentiation medium and medium replaced after 24 hrs.

### EdU incorporation to measure proliferation rate

For over-expression experiments, 54.6-iCEBPB and 16U-iCEBPB myoblasts were seeded in GM with or without 50 ng/mL doxycycline (DOX) (Thermo Fisher Scientific, #J60579). For knock-down experiments human myoblasts lines were treated with either negative control siRNA or anti-CEBPB siRNA as described above. Following 24 hrs in either reagent, medium was replaced with fresh GM, and a 2-hr 5-Ethynyl-2′-deoxyuridine (EdU) pulse was performed using the Click-IT EdU reagent (Invitrogen, #C10337) per manufacturer’s instructions. Cells were then fixed in 4% paraformaldehyde (PFA), permeabilized in PBS/0.1% Triton for 10 min at RT, and incubated in Click-IT EdU AlexaFluor 488 detection reagent (Invitrogen, #C10337) followed by Hoechst33342 (Invitrogen, #C10337) per manufacturer’s instructions. Images were acquired using a Zeiss Axiovert 200M epifluorescence microscope at 10x magnification with a Zeiss AxioCam HRm and AxioVision 4.4 software. The fraction of nuclei that had incorporated EdU was calculated in ImageJ (https://imagej.net/ij/).

### Immunolabeling

For CEBPB over-expression experiments, cells were seeded in either GM or GM + 50 ng/mL DOX. After 48 hrs, GM was removed and changed to either DM or DM + 50 ng/mL DOX. For knock-down experiments, cells were transfected with either 5 ng/mL siNC or 5 ng/mL siCß at the time of seeding and changed to fresh GM after 24 hrs. The following morning, differentiation was induced in DM either containing siNC or siCß and replaced with fresh DM the following day.

After 3-4 days in DM, myotubes were fixed in 4% paraformaldehyde (PFA) for 10 min and permeabilized in PBS/0.1% Triton for 10 min at RT. Cells were blocked in PBS/5% goat serum (Sigma, #G9023) for 1 hr at RT and incubated in primary antibody solution consisting of PBS with 1% goat serum and 1:300 mouse anti-MyHC (DSHB MF-20) and/or 1:300 rabbit anti-CEBPB (Abcam, #ab32358) overnight at 4°C. The following day, secondary antibody incubation was performed for 1 hr at RT in the dark in PBS with 1% goat serum and 1:500 AlexaFluor anti-mouse 488 (ThermoFisher, #A28175) or AlexaFluor anti-rabbit 647 (ThermoFisher, #A21244). Nuclei were stained with 10 µg/mL Hoechst33342 in PBS for 10 min at RT in the dark. Images were acquired using a Zeiss Axiovert 200M epifluorescence microscope with a Zeiss AxioCam HRm and AxioVision 4.4 software at 10x magnification. MyHC-positive area was calculated using a publicly available program (26).

### Examination of CEBPB expression in Muscle RNA-sequencing Data

The Banerji *et al.* (29) FSHD muscle biopsy dataset is available from the European Genome-phenome Archive (EGAS00001007350), and the Wong *et al.* (30) FSHD patient muscle biopsy dataset was downloaded from the authors’ Github repository (https://github.com/FredHutch/RWellstone_FSHD_muscle_biopsy/). Sequencing data from healthy and FSHD myoblasts was obtained from Gene Expression Omnibus (GSE102812). Normalised *CEBPB* expression counts and differential expression analyses were calculated using DESeq2. For DM1 and DMD patient muscle biopsy datasets, *CEBPB* expression was obtained from normalized count matrices accompanying the original publications (DMD: Supplementary Table 2 from Khairallah *et al.* (31) and DM1: (Supplementary Table 5 from Wang *et al.* (32)). *DUX4* over-expression data was downloaded from Jagannathan et al. (33) and Campbell et al. (34), and statistics from the authors’ analyses are reported here.

### Statistical testing

Statistical significance was assessed as described in the figure legends.

## RESULTS

### CEBPB is expressed in human myoblasts and down-regulated upon myogenic differentiation

Translation initiation at three alternative start codons along the CEBPB mRNA transcript yields three distinct protein isoforms: LAP* (44 kDa), LAP (38 kDa) and LIP (20 kDa) (Figure 1A). We first examined the expression dynamics of these three CEBPB isoforms during human myogenesis. To determine if *CEBPB* is expressed in human myoblast cell lines, we analysed our previously published RNA-sequencing data (26). *CEBPB* transcript was expressed at comparable levels in myoblasts from two different control human myoblast lines 54.6 and 16U (Figure 1B), and was significantly down-regulated following *in vitro* differentiation to myotubes in one of the two cell lines (16U) (Figure 1B).

**Figure 1.**
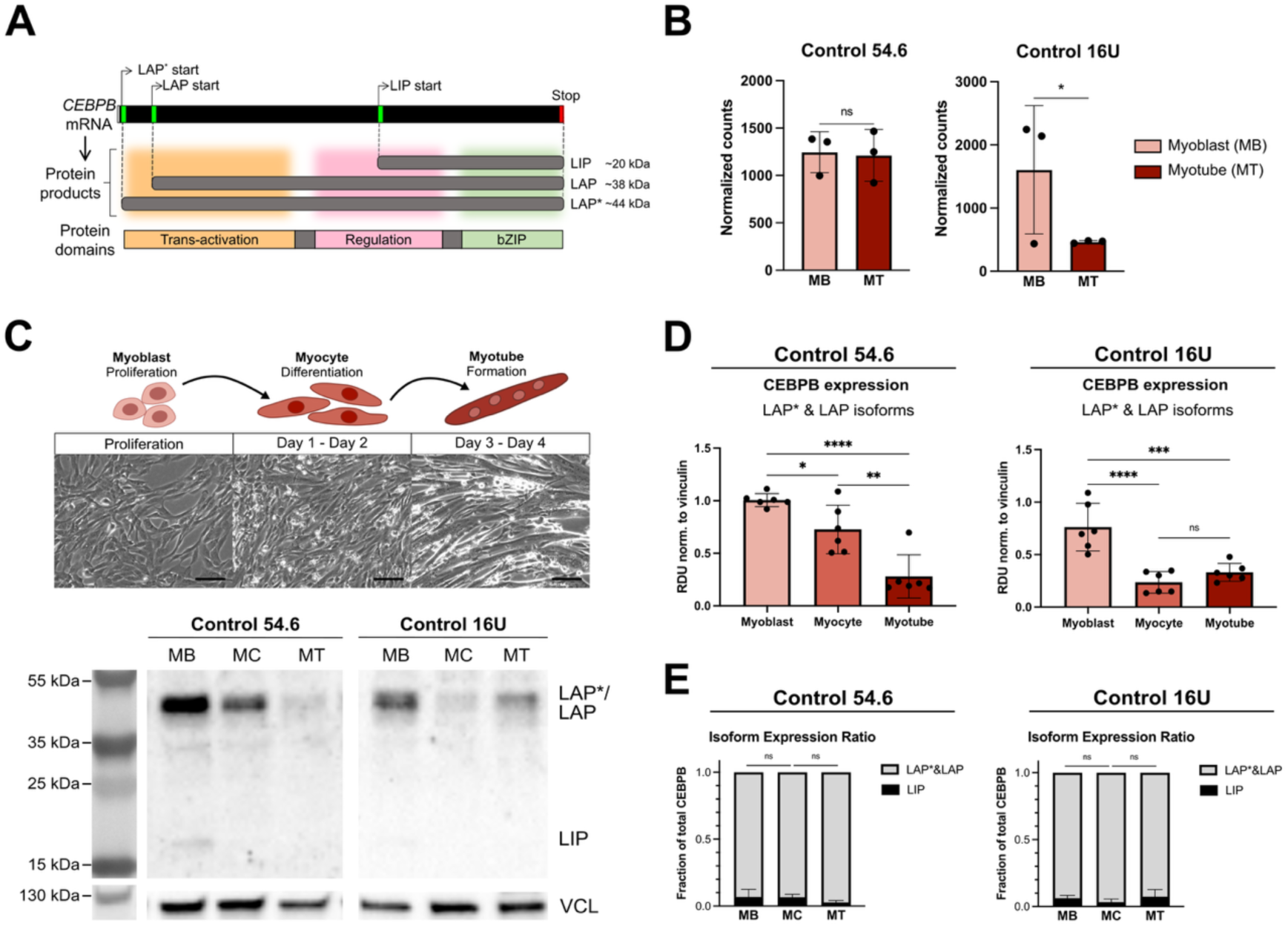
CEBPB protein is present in human myoblasts and down-regulated upon myogenic differentiation. (A) Schematic of CEBPB mRNA transcript and protein isoform generation. Translation initiation at three alternative start codons along the CEBPB mRNA transcript yields distinct protein isoforms: LAP* (44 kDa), LAP (38 kDa) and LIP (20 kDa). The two larger isoforms, LAP* and LAP, contain all three protein domains, while the truncated LIP isoform lacks the trans-activation domain (orange) and part of the regulatory domain (pink). (B) CEBPB mRNA expression levels in publicly available RNA-sequencing data from two human myogenic cell lines cultured as myoblasts (MB) or differentiated into myotubes (MT). Differential expression analysis was performed using DESeq2 with the Wald test and Benjamini-Hochberg correction for multiple testing, where ns denotes *p* > 0.05 and an asterisk denotes *p* < 0.05. (C) Schema and representative brightfield images from each stage of *in vitro* myogenic differentiation examined. Scale bars represent 100 μm. Western blot analysis of CEBPB expression through human myogenesis, with protein levels assessed in myoblasts (MB), myocytes (MC) and myotubes (MT) differentiated from control 54.6 and 16U independent myogenic cell lines. Vinculin (VCL) was used as loading control. (D) Bar plots display quantification of Western blots from (C). Relative density unit (RDU) for LAP*+LAP protein isoform expression normalized to Vinculin (VCL) (top). N= 6 replicates collected over two independent experiments. One-way ANOVA + Tukey post-hoc test, where a single asterisk denotes *p* < 0.05, two asterisks denotes *p* < 0.01, and four asterisks denotes *p* < 0.0001, and ns denotes *p* > 0.05. (E) Stacked bar plots display isoform ratios of LAP*+LAP vs. LIP calculated relative to total CEBPB expression. N= 6 replicates collected over two independent experiments. One-way ANOVA + Tukey post-hoc test, where ns denotes *p* > 0.05.

CEBPB protein levels throughout human myogenesis were also assessed in proliferating myoblasts, differentiating myocytes, and maturing myotubes from each control cell line 54.6 and 16U (Figure 1C). Western blot analysis revealed that LAP*/LAP isoforms, indistinguishable due to their similarity in size, was highest in myoblasts, and significantly reduced in both myocytes and myotubes from both cell lines (Figure 1D). LIP isoform levels were very low compared to LAP*/LAP (Figure 1E) and the ratio of LAP*/LAP to LIP did not significantly change throughout myogenic differentiation (Figure 1E).

### CEBPB regulates human myoblast proliferation

As CEBPB levels were tightly controlled through human myogenesis, we next investigated the functional relevance of this by manipulating *CEBPB* expression. Up-regulation of CEBPB was achieved by generating genetically modified myoblast lines engineered to express *CEBPB* in response to doxycycline (Dox) exposure (Supplementary Figure 1A). Upon Dox administration, both 54.6-iCEBPB and 16U-iCEBPB myoblasts demonstrated robust up-regulation of CEBPB protein by 8 hrs, as detected by immunolabelling (Supplementary Figure 1B) which were the LAP* and LAP isoforms as shown by Western blot (Figure 2A and 2B). Knock-down of CEBPB was achieved using a combination of four anti-CEBPB siRNAs and efficacy verified by Western blot, which demonstrated robust CEBPB knock-down (Figure 2A and 2B).

**Figure 2.**
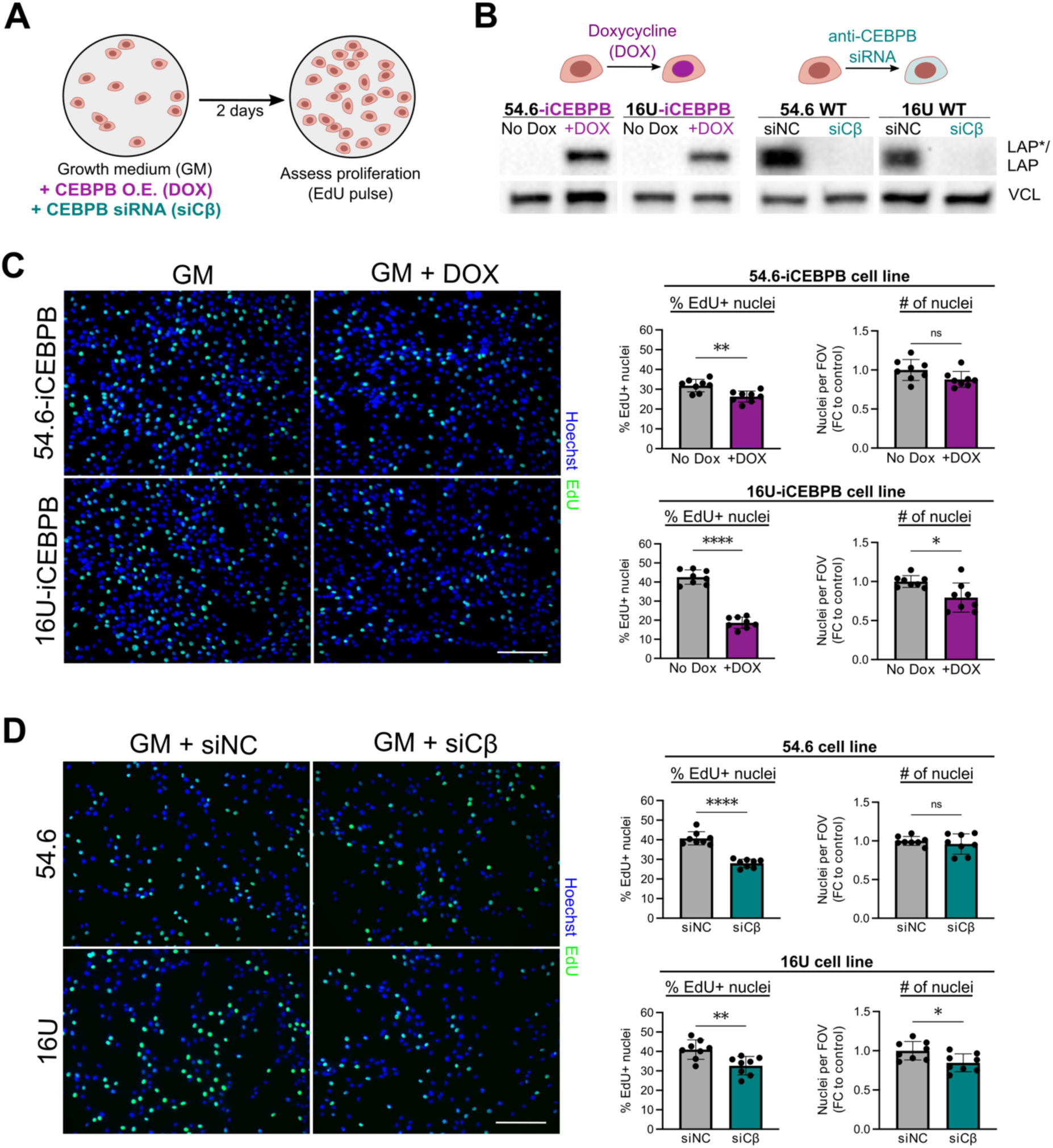
CEBPB regulates human myoblast proliferation. (A) 54.6-iCEBPB and 16U-iCEBPB human myoblast lines were genetically modified to over-express CEBPB (CEBPB O.E.) after exposure to doxycycline (DOX) and were cultured in GM or GM containing DOX for 48 hrs, at which time a 2 hr EdU pulse was performed. Unmodified 54.6 and 16U human myoblast lines were also cultured in GM containing either negative control siRNA (siNC) or anti-CEBPB siRNA (siCβ) overnight, after which growth medium was replaced and EdU pulse performed the following day (2 days post-transfection). (B) Western blots display representative increased CEBPB levels in 54.6-iCEBPB and 16U-iCEBPB cell lines cultured in presence of doxycycline (DOX) for 24 hours (left), and down-regulation of CEBPB in unmodified control 54.6 and 16U myoblasts 48 hours post-transfection with anti-CEBPB (siCβ) siRNA (right), compared to negative control (siNC). Vinculin (VCL) was used as a loading control. (C) Over-expression of CEBPB reduces the myoblast proliferation rate. Quantification of EdU incorporation in 54.6-iCEBPB and 16U-iCEBPB myoblasts following 48 hrs of doxycycline treatment (50 ng/mL). Representative images of incorporated EdU with a Hoechst nuclear counterstain. Bar plots display the percentage of EdU+ nuclei per sample and the total number of nuclei per FOV per sample relative to controls. N=8 replicates per cell line collected over two independent experiments, and each data point represents the average of 3 FOVs per replicate. Total number of nuclei was quantified per sample and divided by the control average of its experiment. Significance was assessed by unpaired t-test, where a single asterisk denotes *p* < 0.05, two asterisks denotes *p* < 0.01, four asterisks denotes *p* < 0.0001, and ns denotes *p* > 0.05. Scale bars represent 200 μm. (D) Decrease in myoblast proliferation rate following knock-down of CEBPB. Quantification of EdU incorporation in control 54.6 and 16U myoblasts 48 hrs post-transfection with either negative control (siNC) or anti-CEBPB (siCβ) siRNA. Representative images of incorporated EdU with a Hoechst nuclear counterstain. Bar plots display the percentage of EdU+ nuclei per sample and the total number of nuclei per FOV per sample relative to controls. N=8 replicates per cell line collected over two independent experiments, and each data point represents the average of 3 FOVs per replicate. Total number of nuclei per sample was divided by the control average of its experiment. Significance was assessed by unpaired t-test, where a single asterisk denotes *p* < 0.05, two asterisks denotes *p* < 0.01, four asterisks denotes *p* < 0.0001, and ns denotes *p* > 0.05. Scale bars represent 200 μm.

To evaluate whether manipulation of CEBPB levels affects the proliferation rate of human myoblasts, EdU incorporation was measured after a two-hour pulse (Figure 2A). Over-expression of *CEBPB* resulted in a significant reduction in EdU incorporation in both control 54.6 and 16U myoblast lines (Figure 2C). siRNA mediated knock-down of *CEBPB* in 54.6 and 16U myoblasts also resulted in decreased EdU incorporation (Figure 2D). Thus, significant divergence from endogenous CEBPB expression levels negatively impacts the proliferation rate of human myoblasts. Importantly, there was no evidence of myotube formation, indicating that myoblasts were not precociously fusing in growth culture conditions, unlike what has previously been reported for murine myoblasts (17).

### CEBPB regulates human myoblast differentiation

Next, we assessed whether manipulation of *CEBPB* expression impacts human myogenic differentiation and fusion. We either knocked down or over-expressed CEBPB expression during either myoblast proliferation, differentiation, or both proliferation and differentiation, and evaluated myotube formation (Figure 3A). Non-induced 54.6-iCEBPB and 16U-iCEBPB myoblast lines both formed substantial numbers of myotubes following 3-4 days in differentiation medium (DM), as revealed by immunolabelling for MyHC (Figure 3B). However, when proliferating 54.6-iCEBPB or 16U-iCEBPB myoblasts were treated with doxycycline (to up-regulate CEBPB levels), myogenic differentiation and myotube formation were dramatically impaired in both lines (Figure 3B). Up-regulation of CEBPB levels using doxycycline at the onset of myogenic differentiation resulted in nuclear-located CEBPB, as revealed by immunolabelling, which again blocked nearly all myogenic differentiation and myotube formation (Figure 3B).

**Figure 3.**
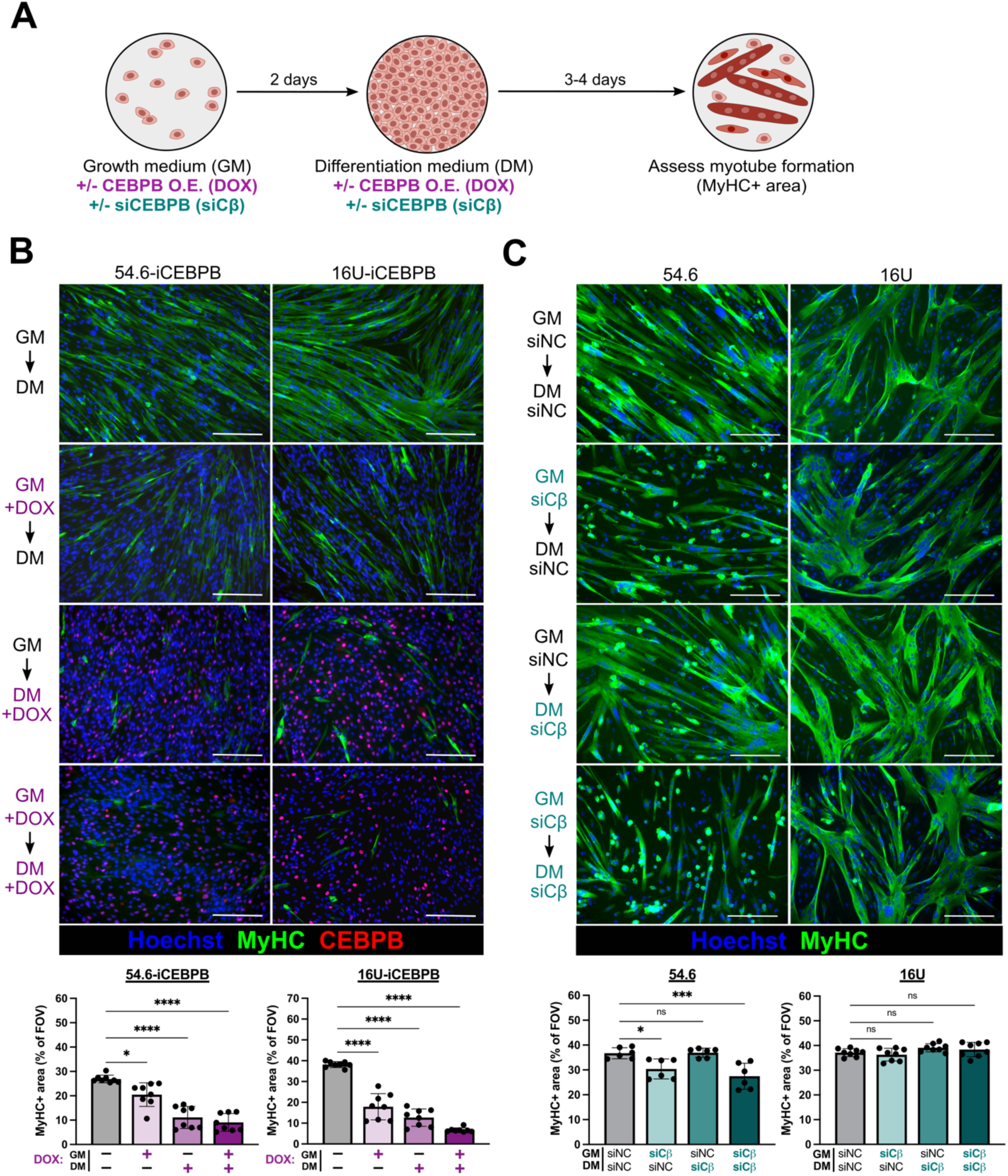
Up-regulating CEBPB inhibits myogenic differentiation. (A) Experimental overview: proliferating human myoblasts were grown in growth medium (GM) until confluency, then switched to differentiation medium (DM). Myotube formation was assessed by quantification of the area per field occupied by myosin heavy chain (MyHC). Over-expression (CEBPB O.E.) or knock-down (siCEBPB) of CEBPB was performed either during proliferation only, during differentiation only, or during both proliferation and differentiation. (B) Up-regulation of CEBPB prevents myogenic differentiation. Quantification of MyHC+ area shows myotube formation of 54.6-iCEBPB or 16U-iCEBPB is impaired moderately with addition of doxycycline during proliferation (GM+DOX → DM), and dramatically if doxycycline is added at the onset of differentiation (GM → DM+DOX and GM+DOX → DM+DOX). Representative images immunolabelled for MyHC and CEBPB, and counterstained with Hoechst. One-way ANOVA + Tukey post-hoc test, where a single asterisk denotes *p* < 0.05 and four asterisks denotes *p* < 0.0001. N=8 replicates per cell line collected over two independent experiments, with data point representing the mean MyHC+ area across 3 FOVs per replicate. Scale bars represent 200 μm. (C) Reduction of CEBPB has little effect on myogenic differentiation. Myotube formation was assessed in control 54.6 and 16U in the presence of anti-CEBPB siRNA (siCβ) during myoblast proliferation and/or the onset of myogenic differentiation. Quantification of MyHC+ area revealed that reducing CEBPB levels during proliferation (GM+siCβ ◊ DM+siNC) mildly impaired differentiation in one line, while decreasing CEBPB during differentiation (GM+siNC → DM+siCβ and GM+siCβ ◊ DM+siCβ) had no effect. Representative images immunolabelled for MyHC. One-way ANOVA + Tukey post-hoc test, where a single asterisk denotes *p* < 0.05, four asterisks denotes *p* < 0.0001, and ns denotes *p* > 0.05. N=6-8 replicates per cell line collected over two independent experiments, with data point representing the mean MyHC+ area across 3 FOVs per replicate. Scale bars represent 200 μm.

The impact on myogenic differentiation of reduced *CEBPB* expression during either proliferation, differentiation, or both proliferation and differentiation, was then assessed in 54.6 and 16U myoblasts (Figure 3A). siRNA-mediated knock-down of *CEBPB* expression during myoblast proliferation or both myoblast proliferation and differentiation had a minor negative effect on subsequent differentiation/myotube formation in control 54.6, but not control 16U (Figure 3C). Knock-down of *CEBPB* during differentiation alone did not affect myogenic differentiation or myotube formation in either cell line (Figure 3C), consistent with the observation that CEBPB is normally down-regulated following the onset of myogenic differentiation (Figure 1D). Combined, these results establish the expression dynamics of *CEBPB* during human myogenesis and demonstrate that reduction of CEBPB is required for effective myotube formation.

### CEBPB expression is altered in muscular dystrophy

Characterization and manipulation of CEBPB revealed that CEBPB expression is tightly controlled during healthy human myogenesis, and that deviation from normal expression dynamics has deleterious effects on proliferation and differentiation in human myoblasts. Given these observations, we next examined whether alterations in *CEBPB* expression were relevant in the context of muscle disease, using publicly available data from Duchenne muscular dystrophy (31) (Figure 4A), myotonic dystrophy type 1 (32) (Figure 4B) or FSHD (29, 30) (Figure 4C, 4D).

**Figure 4.**
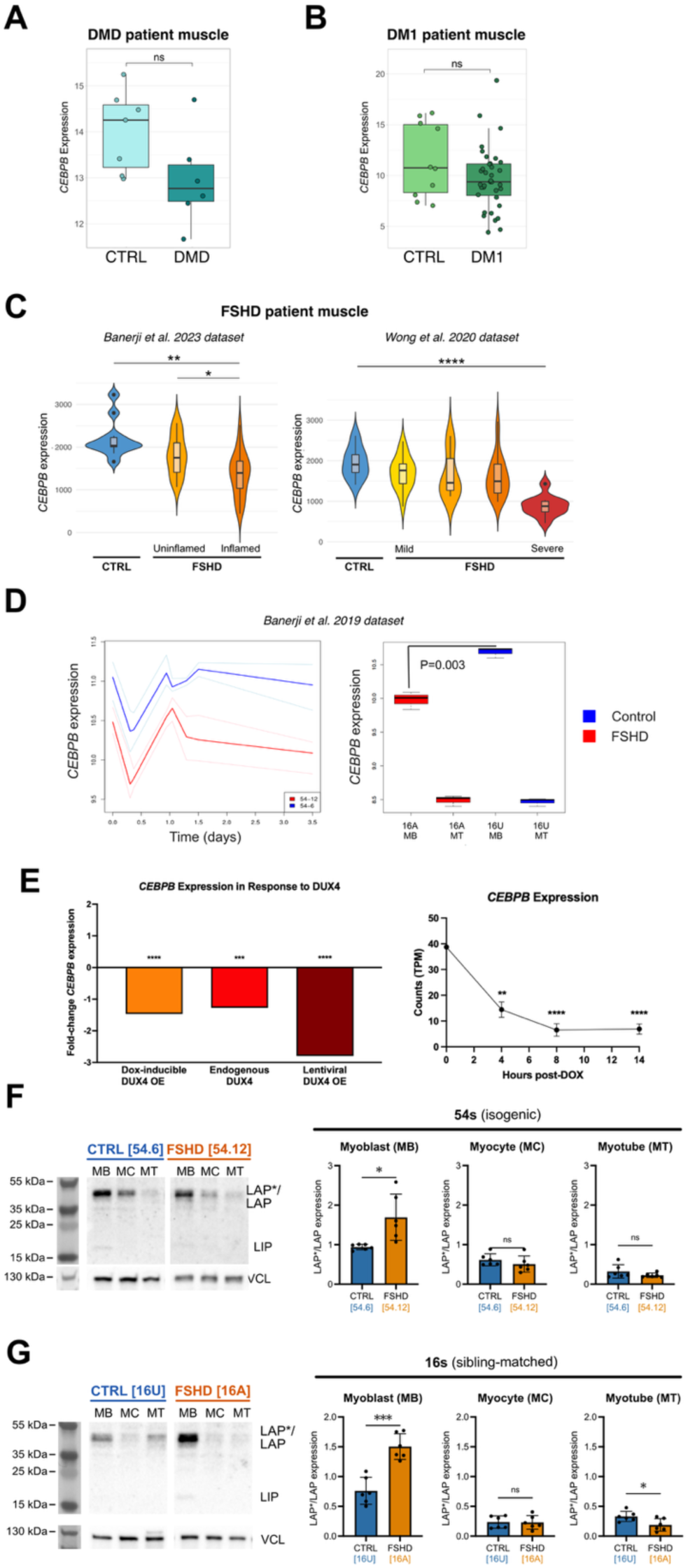
Mis-regulation of CEBPB in FSHD muscle. (A) Box plot displays expression of *CEBPB* in bulk transcriptomic data from muscle biopsies from 6 healthy control individuals (light blue) and 7 patients with DMD (dark blue) reported by Khairallah *et al.* (31). Normalized counts were downloaded from the original publication’s supplementary data (31). Boxes represent the interquartile range (IQR), lines display the median, and whiskers depict the smallest and largest values within 1.5*IQR from the IQR. Statistical significance was assessed by Wilcoxon ranked-sum test, where ns denotes *p* > 0.05. Box plot displays expression of *CEBPB* in bulk transcriptomic data from muscle biopsies from 10 healthy control individuals (light green) and 36 patients with DM1 (dark green) reported by Wang *et al.* (32). Normalized counts were downloaded from the original publication’s supplementary data (32). Boxes represent the interquartile range (IQR), lines display the median, and whiskers depict the smallest and largest values within 1.5*IQR from the IQR. Statistical significance was assessed by Wilcoxon ranked-sum test, where ns denotes *p* > 0.05. (B) Violin plots display expression of *CEBPB* in Banerji *et al.* 2023 (29) and Wong *et al.* (30) datasets. Differential expression analysis was performed using DESeq2 with the Wald test and Benjamini-Hochberg correction for multiple testing, where a single asterisk denotes *p* > 0.05, two asterisks denotes *p* < 0.01, and four asterisks denotes *p* < 0.0001. All analyses were unpaired, with the exception of TIRM^−^ vs. TIRM^+^ samples in Banerji dataset, in which a paired analysis was performed between matched samples from individual patients. Boxes represent the interquartile range (IQR), lines display the median, and whiskers depict the smallest and largest values within 1.5*IQR from the IQR. (C) Expression of *CEBPB* in RNA-sequencing data from 54.6 control cell line (blue) and 54.12 FSHD myoblast cell line (red), as well as 16U control cell line (blue) and 16A FSHD cell line (red) at various time points throughout *in vitro* myogenesis (26). The ‘54s’ cell line pair was isolated from a mosaic patient, with 54.6 (blue) derived from a clone containing a healthy genetic background and 54.12 (red) containing the disease-conferring contraction of the D4Z4 region. The ‘16s’ cell lines were derived from an FSHD patient (‘16A’, red) and a sibling matched control (‘16U’, blue). (D) Decrease in *CEBPB* expression in response to endogenous and over-expressed DUX4, which begins as soon as 4 hours after *DUX4* induction. Log fold-changes and RNA-sequencing data, along with their corresponding statistics, were obtained from the Supplementary materials from Jagannathan et al. (33) or Campbell et al. (34). (F, G) CEBPB protein expression was compared in two pairs of control vs. FSHD immortalized myoblast cell lines. Each pair was differentiated in parallel, and protein samples were collected from proliferating myoblasts (MB), from differentiating myocytes (MC) 24 hours after induction of differentiation, and from myotubes (MT). CEBPB protein expression was evaluated by Western blot, and levels of LAP*/LAP isoforms were quantified relative to vinculin (VCL). Colorimetric image of pre-stained protein ladder shown on right to estimate molecular weights. N=6 replicates per cell line collected from two independent experiments. Statistical significance was assessed by unpaired t-test, with one asterisk denoting *p* < 0.05, three asterisks denoting *p* < 0.001, and ns denoting *p* > 0.05.

Expression of *CEBPB* in RNA-sequencing data from skeletal muscle biopsies of patients with Duchenne muscular dystrophy (Figure 4A) or myotonic dystrophy type 1 (Figure 4B) was not significantly altered compared to healthy control individuals. However, in RNA-sequencing data of muscle biopsies from two independent cohorts of FSHD patients, *CEBPB* expression was significantly decreased compared to healthy controls, which was correlated with increasing disease severity in both datasets (Figure 4C). As studies using bulk muscle tissue samples encompass gene expression from multiple cell types, to specifically investigate *CEBPB* expression in myogenic cells from FSHD, we next examined a time series RNA-sequencing dataset collected from unaffected control and FSHD myoblasts undergoing myogenic differentiation (Figure 4D) (26). Consistent with bulk muscle biopsy data (Figure 4C), levels of *CEBPB* mRNA were also lower in the FSHD myoblasts across all time points compared to controls (Figure 4D).

The two myoblast lines that were used to characterize the role of CEBPB in healthy human myogenesis both have FSHD counterparts. The ‘54’ myoblast lines were expanded from a mosaic male FSHD1 patient to generate clones with both a normal genetic background (e.g. 54.6) or myoblast lines containing the D4Z4 contraction (e.g. 54.12), and so are isogenic with the exception of the D4Z4 contraction (27). The ‘16U’ myoblast line was isolated from an unaffected woman, while 16A was derived from her sister with FSHD1 (28).

We further confirmed this transcriptional down-regulation of *CEBPB* by examining publicly available transcriptomic data (33) from a study comparing three different *in vitro* models of DUX4 over-expression in human myoblasts (lentiviral transduction, doxycycline-induced, and endogenous levels in FSHD patient-derived cells), and observed that *CEBPB* expression was consistently down-regulated across all three models (Figure 4E). Analysis of a separate publicly available dataset reported by the same group (34), showed that significant down-regulation of *CEBPB* occurs as early as 4 hours following DUX4-overexpression (Figure 4E).

CEBPB protein levels dropped during myogenic differentiation and fusion into myotubes in FSHD 54.12 and 16A cell lines (Figure 4F and G), as observed in 54.6 and 16U control lines. In contrast to the decreased *CEBPB* gene expression observed both in FSHD patient biopsies and cell lines (Figure 4C and D), CEBPB protein levels were significantly higher in proliferating FSHD 54.12 or 16A myoblasts compared to control myoblasts (Figure 4F and G). This finding was interesting considering the negative effect of abnormal CEBPB levels on human myogenic differentiation (Figure 3), given that some FSHD myoblast clones differentiate poorly (26). CEBPB protein levels in FSHD and control myocytes were not significantly different in either cell line pair (Figure 4F and G). In myotubes, CEBPB protein expression was unaltered between FSHD 54.12 and control 54.6 (Figure 4F) but was slightly lower in FSHD 16A myotubes than control 16U (Figure 4G).

### CEBPB down-regulation improves myogenic differentiation in FSHD

To evaluate whether decreasing CEBPB protein levels could restore myotube formation in FSHD cells, 54.12 and 16A myoblasts during proliferation and/or at the onset of differentiation were transfected with anti-CEBPB siRNA (siCβ) or negative control siRNA (siNC). As expected, FSHD myotubes transfected with control siNC exhibited impaired myogenesis compared to their respective controls with siNC (Figure 5A and B).

**Figure 5:**
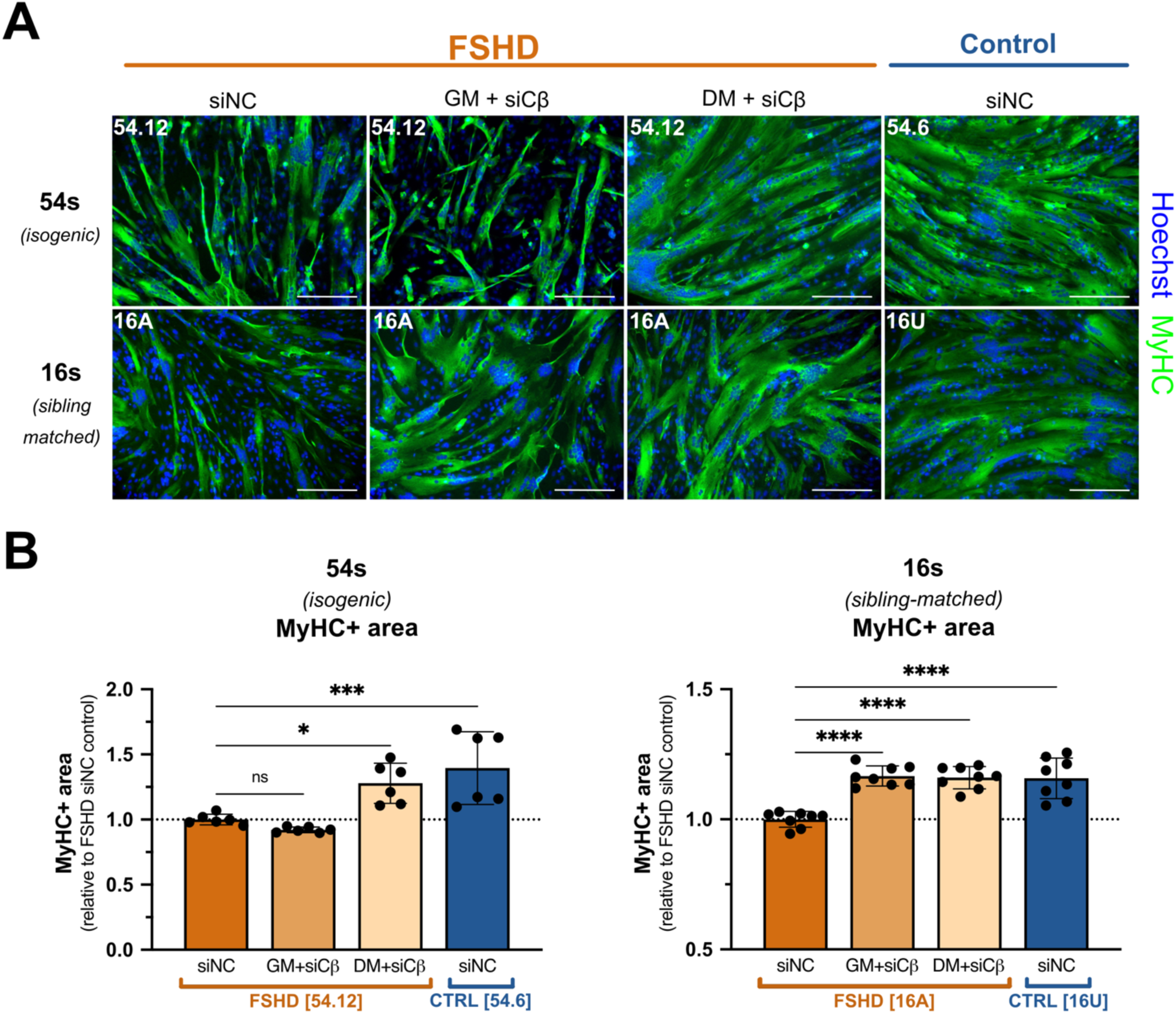
Reducing CEBPB levels improves myogenic differentiation in FSHD. (A) FSHD cell lines 54.12 and 16A were differentiated with addition of negative control siRNA (siNC) or anti-CEBPB siRNA (siCβ) during either myoblast proliferation (in growth medium; GM) or at the onset of differentiation (in differentiation medium; DM). Representative images of myotubes immunolabelled for MyHC and counterstained with Hoechst. As expected, FSHD myotubes exhibit impaired myogenesis compared to their respective control cell lines 54.6 or 16U (siNC conditions). CEBPB knock-down in FSHD myoblasts during differentiation improved myotube formation to similar levels to that in their respective control clones 54.6 or 16U. Scale bars represent 200 μm. (B) Quantification of MyHC+ area in each cell line and condition. N = 6-8 replicates collected over 2 independent experiments. Significance was assessed by one-way ANOVA with Tukey’s post-hoc test, where one asterisk denotes *p* < 0.05, three asterisks denotes *p* < 0.001, four asterisks denotes *p* < 0.0001, and ns denotes *p* > 0.05.

While we found that knock-down of CEBPB in proliferating myoblasts mildly impaired differentiation of the control 54.6 myoblast line (Figure 3C), it had no effect on myogenic differentiation in the FSHD 54.12 line (Figure 5A and B). However, CEBPB knock-down in proliferating 16A FSHD myoblasts enhanced differentiation (Figure 5A and B), an effect which was not observed in their control 16U counterpart (Figure 3C). Furthermore, knock-down of CEBPB at the onset of myogenic differentiation improved differentiation of both FSHD 54.12 and 16A myoblast lines to similar levels to their respective controls transfected with siNC (Figure 5A and B). These results show that elevated CEBPB levels in FSHD myogenic cells may contribute to impaired differentiation/myotube formation in FSHD.

## Discussion

Our aim was to characterise the role of CEBPB in human myogenesis and evaluate its potential contribution to FSHD. Previous studies in mice have shown that CEBPB is expressed in quiescent satellite cells and macrophages within skeletal muscle (14, 17–19).

Investigating CEBPB protein level in human myoblasts cultured as proliferating myoblasts, differentiating myocytes or differentiated myotubes revealed that CEBPB protein levels decrease through differentiation. This is consistent with studies in mice demonstrating rapid down-regulation of CEBPB mRNA and protein at the onset of myogenic differentiation (17). LAP* and LAP were the predominant CEBPB isoforms present in human myoblasts, accounting for >90% total CEBPB content. This ratio does not change significantly over the course of myogenic differentiation.

Inducible up-regulation of the full-length CEBPB transcript containing all three alternative start codons in genetically engineered human control 54.6-iCEBPB and 16U-iCEBPB myoblasts showed that the LAP*/LAP isoforms, but not LIP, were dramatically up-regulated. Although CEBPB expression is highest in myoblasts, either up-regulation or knock-down of CEBPB reduced the myoblast proliferation rate. Reduction of proliferation following CEBPB over-expression may be explained by the fact that CEBPB promotes quiescence, and is a direct transcriptional activator of *Pax7* (18). Genetic ablation of *CEBPB* in murine satellite cells using a *Cebpb^fl/fl^Pax7^CreER/+^* mouse model causes increased proliferation in primary satellite cell-derived myoblasts (18). In contrast, we observed reduced proliferation following knock-down of CEBPB in immortalized human myoblasts. This difference may be due to the stage at which CEBPB function was inactivated. Our knock-down was performed in proliferating human myoblasts, a more advanced stage in myogenic progression compared to the mouse model (18), where CEBPB function was inactivated from quiescent/activated murine satellite cells before proliferation. Loss of CEBPB in murine satellite cells also up-regulates genes important for the later stages of regeneration such as myogenin and promotes myogenic differentiation (17), and myoblasts must withdraw from cell cycle to differentiate (35, 36). Thus, our knock-down of CEBPB in human myoblasts may have removed repression of genes important for myogenic progression and so signaled cell cycle arrest prior to differentiation. However, this was not accompanied by any obvious formation of myotubes.

In addition to regulating proliferation, we also found that down-regulation of CEBPB is an important pre-requisite for myogenic differentiation in human. Up-regulation of CEBPB in either proliferating human 54.6-iCEBPB or 16U-iCEBPB myoblasts led to compromised differentiation and significantly fewer and smaller myotubes. However, up-regulation of CEBPB during differentiation abrogated myotube formation. This is consistent with observations that viral over-expression of CEBPB in both primary murine myoblasts, as well as in proliferating murine C2C12 cells, impairs subsequent myogenic differentiation (17).

There was no significant improvement in myotube formation in human myoblasts following siRNA knock-down of CEBPB during differentiation. This contrasts with data from mouse, in which myogenic differentiation was improved when CEBPB levels were reduced, as shown by increased fusion of primary satellite cells isolated from *Cebpb^fl/fl^Pax7^CreER/+^* mice following tamoxifen induction and muscle fiber hypertrophy *in vivo* (17). Knock-down of CEBPB in proliferating myoblasts did, however, have a negative effect on subsequent myogenesis in one control line.

Given the importance of CEBPB as a regulator of both murine and human myogenesis, it is reasonable to examine whether it is perturbed in disease states. We examined *CEBPB* expression in three different types of muscular dystrophy: FSHD, Duchenne muscular dystrophy and myotonic dystrophy type 1. Only RNA-sequencing data from muscle biopsies of FSHD patients (29, 30, 37) showed a difference: a significant reduction of *CEBPB* compared to controls.

We further examined mis-regulation of *CEBPB* in FSHD using immortalized human myoblasts derived from FSHD patients, comparing *CEBPB* expression dynamics in healthy 54.6 and 16U myoblast lines to their FSHD counterparts (54.12 and 16A). RNA-sequencing data from a time-course comparison of the control 54.6 cell line versus the FSHD 54.12 cell line throughout myogenic differentiation suggested that *CEBPB* mRNA levels are consistently reduced in FSHD myogenic cells. We confirmed that DUX4 directly leads to transcriptional down-regulation of *CEBPB* using publicly available transcriptomic data (33), and that significant down-regulation of *CEBPB* occurs as early as 4 hours following *DUX4* induction (34).

As in healthy human myogenesis, CEBPB protein levels were highest in proliferating FSHD myoblasts compared to myocytes and myotubes. However, unlike that observed at the transcriptional level, CEBPB protein levels in proliferating FSHD myoblasts were consistently elevated compared to healthy controls. RNA and protein levels do not always correlate (38), and our observations could be due to several non-mutually exclusive reasons, including post-transcriptional and post-translational regulation, cellular localization, and differences in mRNA and protein stability. CEBPB mRNA half-life is approximately 40-60 minutes in adipocytes and lymphomas, however it can be stabilized by the RNA-binding protein human antigen R (HuR), which binds the 3’ UTR and extends half-life for up to 2 hours (39, 40). While stabilization of CEBPB mRNA has not been reported in muscle, HuR promotes myogenesis through stabilization of other myogenic regulators such as MyoD and Myogenin (41), and is a putative DUX4 binding partner (42). If HuR-mediated stabilization of CEBPB transcripts is disrupted in FSHD through interaction with DUX4, this may explain the reduced transcript levels in FSHD patients, and is consistent with CEBPB expression being down-regulated by DUX4 (33, 34).

The elevated protein levels may also be due to differences in CEBPB protein stability in FSHD. In murine C2C12 myoblasts, Mouse double minute 2 (Mdm2) ubiquitin ligase targets CEBPB for degradation, and knock-down of Mdm2 results in high CEBPB levels and a blockade in myogenic differentiation (43). MDM2 can be linked to FSHD through its inhibitory effect on p53 (44), a pathway which has been associated with both DUX4-induced myopathy (45) as well as *DUX4* expression in the early embryo and in FSHD cell lines (46), although other studies suggest DUX4 cytotoxicity is independent of the p53 axis (47). While the relationship between DUX4 and the p53 pathway remains under debate, any alterations in p53 activity in FSHD could influence degradation of other MDM2 targets such as CEBPB, potentially preventing their degradation and leading to the accumulation of CEBPB protein observed in FSHD myoblasts.

Interestingly, the compromised differentiation caused by up-regulation of CEBPB in proliferating human control myoblasts prior to the onset of differentiation is a similar phenotype to that observed in some FSHD clones, which form hypotrophic myotubes (26). Knocking down CEBPB in FSHD myoblasts either during proliferation or at the onset of differentiation allowed assessment of whether elevated CEBPB levels contributes to impaired differentiation in FSHD. Reduction of CEBPB at the start of differentiation led to significant improvement in myotube formation in both FSHD myoblast lines examined, while knockdown during proliferation (when CEBPB levels are highest) only rescued differentiation in the FSHD 16A line. This is interesting given that knock-down of CEBPB at the onset of differentiation did not induce hypertrophy in the unaffected control cell lines, yet was able to rescue defective FSHD myogenesis.

Our data suggest that excessive accumulation, or improper clearance, of CEBPB during myogenic differentiation impairs subsequent myogenesis. However, by 24 hours following induction of differentiation, CEBPB expression in FSHD myocytes resolved to that of control levels, meaning the consequence of this accumulation occurs early following the onset of differentiation. The sequence and timing of gene expression changes during myogenesis are tightly and temporally controlled, and CEBPB is a transcriptional regulator of many of these genes (17, 18). Thus, these findings suggest that transcriptional control maintained by CEBPB needs to be lifted at a specific time, and if this window is missed then excess CEBPB is present at the onset of myogenesis that will impair differentiation, as is the case for FSHD. This would explain why myogenesis is still defective in FSHD cells after 24 hours even though protein levels are reduced to similar levels as controls by this time.

## Conclusion

In summary, CEBPB is a regulator of human myogenesis, is expressed in human myoblasts, and is down-regulated upon myogenic differentiation. Reduction of CEBPB protein level is necessary for myogenic differentiation and fusion to proceed. In FSHD, CEBPB expression dynamics are mis-regulated, and increased CEBPB protein levels in FSHD myoblasts may contribute to defective myogenesis in FSHD.

## Supporting information

Engquist et al CEBPB biorxiv 2025- Supplementary Figure 1

## Acknowledgements

We would like to thank: Vincent Mouly (Center for Research In Myology, UMRS 974 Sorbonne Université-INSERM, Paris, France) and Charles P. Emerson (Wellstone Muscular Dystrophy Program, University of Massachusetts Medical School, MA, USA) for providing immortalised myoblast lines; ENE was funded by Wellcome Trust PhD Studentship (WT 222352/Z/21/Z) and SOLVE FSHD. The Zammit laboratory is funded by the Medical Research Council (MR/P023215/1 and MR/S002472/1), the FSHD Society, Friends of FSH Research grant (FSHS-82013-06) and Association Française contre les Myopathies. CRSB was supported by the CRUK City of London Centre Award [CTRQQR-2021\100004].

