## Supplementary material for "CEBPβ regulates myoblast proliferation and myogenic differentiation during human myogenesis and rescues defective differentiation in FSHD": Engquist et al CEBPB biorxiv 2025- Supplementary Figure 1

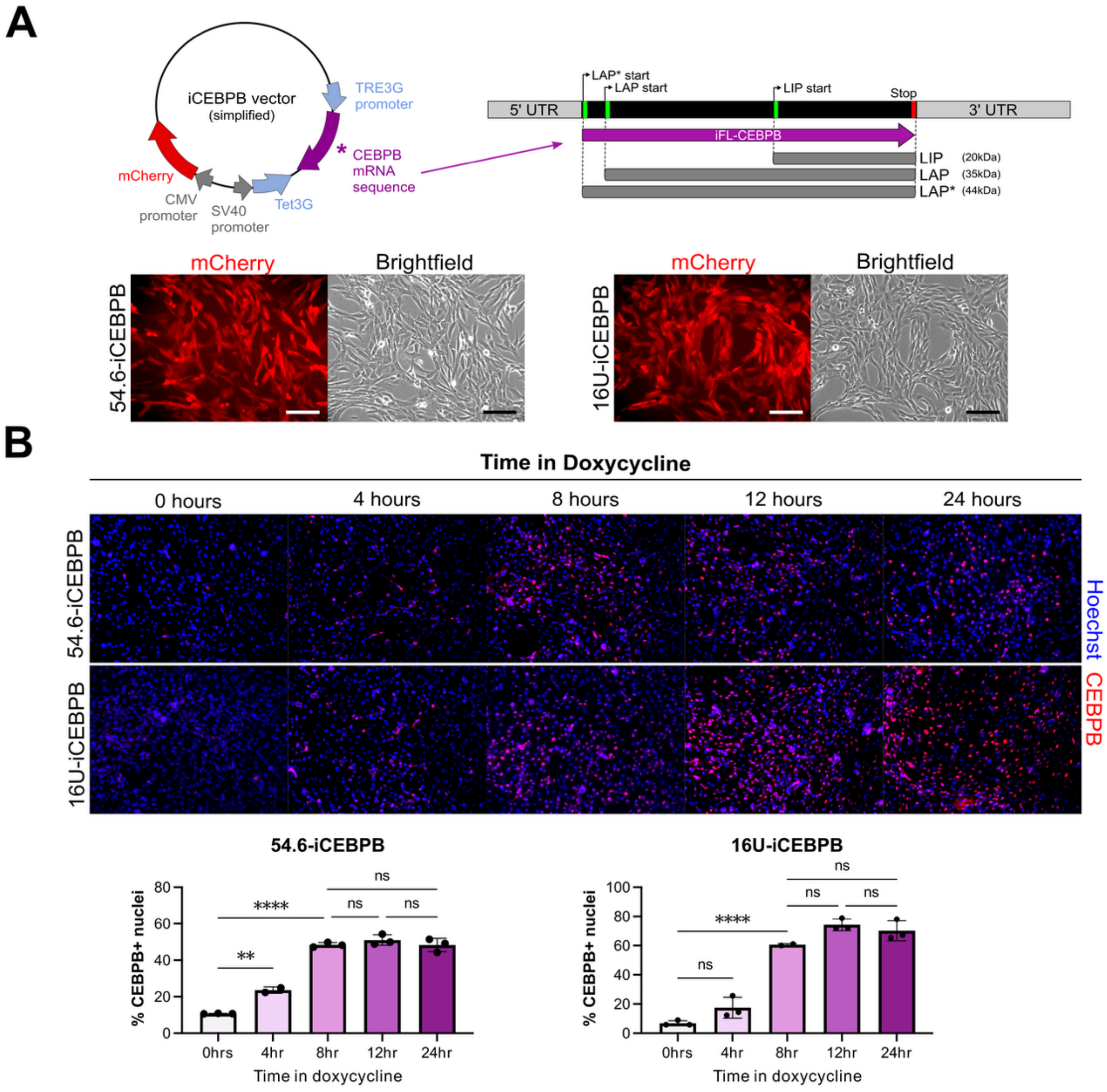


**Supplementary Figure 1. Generation of genetically modified cell lines with doxycycline-inducible over- expression of CEBPB.**

(A) Simplified schematic of lentiviral vector used to generate 54.6-iCEBPB and 16U-iCEBPB cell lines. The full-length *CEBPB* coding sequence is driven by the *TRE* promoter, and ubiquitously active *SV40* and *CMV* promoters respectively drive expression of the reverse tetracycline-controlled transactivator and *mCherry*. Representative images of 54.6-iCEBPB and 16U-iCEBPB cell lines after FACs sorting for mCherry+ cells, resulting in purified populations of cells with successful integration of the vector. Scale bars represent 200 µm.

(B) Time course of CEBPB over-expression following doxycycline treatment. 54.6-iCEBPB (top) and 16U (bottom) myoblasts were induced with doxycycline for either 0 hours (non-induced control), 4 hrs, 8 hrs, 12 hrs, or 24 hrs prior to fixation. Top panel displays representative immunofluorescence images of cells labelled for CEBPB (red) and counterstained with Hoechst33342 (blue) to highlight all nuclei. Bar graphs below display the percentage of nuclei expressing CEBPB at each time point. N=3 replicates per cell line per time point, and each data point represents the mean % CEBPB+ cells from 3 FOVs per replicate. Statistical significance was assessed by one-way ANOVA with Tukey's post-hoc test for multiple comparisons, where two asterisks denotes $p < 0.01$, four asterisks denote $p < 0.0001$, and ns denotes $p > 0.05$.
